# Insulator proteins contribute to expression of gene loci repositioned into heterochromatin in the course of *Drosophila* evolution

**DOI:** 10.1101/802371

**Authors:** Sergei Yu. Funikov, Alexander P. Rezvykh, Dina A. Kulikova, Elena S. Zelentsova, Lyubov N. Chuvakova, Venera I. Tyukmaeva, Irina R. Arkhipova, Michael B. Evgen’ev

**Affiliations:** Engelhardt Institute of Molecular Biology of Russian Academy of Sciences, Moscow, Russia; Moscow Institute of Physics and Technology, Dolgoprudny, Moscow Region, Russia; Koltzov Institute of Developmental Biology of Russian Academy of Sciences, Moscow, Russia; Department of Biological and Environmental Science, University of Jyväskylä, 40014, Finland; Josephine Bay Paul Center for Comparative Molecular Biology and Evolution, Marine Biological Laboratory, Woods Hole, Massachusetts, USA

**Keywords:** Drosophila, heterochromatin, gene expression, molecular evolution, insulators, BEAF-32

## Abstract

Pericentric heterochromatin in *Drosophila* is generally composed of repetitive DNA forming a transcriptionally repressive environment. Nevertheless, dozens of genes were embedded into pericentric genome regions during evolution of *Drosophilidae* lineage and retained functional activity. However, factors that contribute to “immunity” of these gene loci to transcriptional silencing remain unknown. Here, we investigated molecular evolution of the essential *Myb* and *Ranbp16* genes. These protein-coding genes reside in euchromatic loci of chromosome X in *D. melanogaster* and related species, while in other studied *Drosophila* species, including evolutionary distant ones, they are located in genomic regions highly enriched with the remnants of transposable elements (TEs), suggesting their heterochromatic nature and location. The promoter region of *Myb* exhibits a conserved structure throughout the *Drosophila* phylogeny and carries motifs for binding of chromatin remodeling factors, including insulator BEAF-32, regardless of eu- or heterochromatic surroundings. Importantly, BEAF-32 occupies not only the promoter region of *Myb* but is also found in the vicinity of transcriptional start sites (TSS) of *Ranbp16* gene as well as in a wide range of genes located in the contrasting chromatin types in *D. melanogaster* and *D. virilis,* denoting the boundary of the nucleosome-free region available for RNA polymerase II recruitment and the surrounding heterochromatin. We also find that along with BEAF-32, insulators dCTCF and GAF are enriched at the TSS of heterochromatic genes in *D. melanogaster*. Thus, we propose that insulator proteins contribute to gene expression in the heterochromatic environment and, hence, facilitate the evolutionary repositioning of gene loci into heterochromatin.

**Author summary:** Heterochromatin in *Drosophila* is generally associated with transcriptional silencing. Nevertheless, hundreds of essential genes have been identified in the pericentric heterochromatin of *Drosophila melanogaster*. Interestingly, genes embedded in pericentric heterochromatin of *D. melanogaster* may occupy different genomic loci, euchromatic or heterochromatic, due to repositioning in the course of evolution of *Drosophila* species. By surveying factors that contribute to the normal functioning of the relocated genes in distant *Drosophila* species, i.e. *D. melanogaster* and *D. virilis*, we identify certain insulator proteins (e.g.BEAF-32) that facilitate the expression of heterochromatic genes in spite of the repressive environment.

## Introduction

Eukaryotic genomes are packaged in chromatin consisting of DNA and associated proteins. Typically, chromatin can be divided into two basic forms, euchromatin and heterochromatin (1). These types of chromatin are distinguished by several distinctive properties, including DNA sequence composition, specific histone modifications and binding proteins, nuclear and chromosomal localization, and frequency of meiotic recombination (1, 2). One of the major subtypes of heterochromatin in *Drosophila* is marked by heterochromatin protein 1 (HP1a) and di- or trimethylated H3K9 (3, 4). This subtype of heterochromatin covers large genomic segments primarily around centromeres and, in association with the protein POF (painting of fourth), the entire dot chromosome 4 in *D. melanogaster* (3–5). Pericentric heterochromatin is mainly composed of repetitive sequences, including remnants of various transposable elements (TEs), satellite DNAs and other repeats (6). A distinctive feature of heterochromatin is the ability to silence euchromatic genes placed within heterochromatic environment due to chromosomal inversions or transposition events, a phenomenon called position effect variegation (PEV) (7–12). Transcriptional silencing of euchromatic genes in PEV is mediated by spreading of heterochromatin-associated marks HP1a and H3K9me3 across the gene loci transferred to heterochromatin (8, 10).

Despite the repressive environment, dozens of essential genes were identified in the pericentric heterochromatin of *D. melanogaster* (13–16). Interestingly, genes embedded in pericentric heterochromatin in *D. melanogaster* may occupy distinct genomic loci, euchromatic and heterochromatic, in other *Drosophila* species (17). For instance, two adjacent genes *RpL15* and *Dbp80* located in the pericentric region of chromosome 3L in *D. melanogaster* reside in a euchromatic region in *D. pseudoobscura* (18). A similar pattern was demonstrated for genes *light* and *Yeti* located in pericentric regions in *D. melanogaster,* while in *D. virilis* they are found within euchromatin on the same chromosomal elements (19, 20). Recently, it was shown that most of the pericentric genes found at both arms of chromosome 2 of *D. melanogaster* are located in euchromatic loci in the *D. virilis* genome (21).

However, although repositioning of genes between euchromatin and heterochromatin during genome evolution is not unusual in the *Drosophilidae* lineage, the “immunity” of heterochromatic genes to the transcriptionally repressive environment remains paradoxical and unexplained. Thus, it is not clear whether these loci have undergone adaptation to heterochromatic environment or had some intrinsic properties permitting local adaptation. Previously, it was shown that molecular organization of promoter regions is largely conserved between heterochromatic and euchromatic genes, indicating that adaptation to heterochromatin probably does not require major changes in regulatory sequences (20). However, expression of heterochromatic genes requires the methylated H3K9 mark (22, 23), and the ability to repositioning during evolution is a characteristic feature of gene clusters that show close association with HP1a protein (21).

Chromatin insulator elements and associated proteins were originally defined by their ability to protect transgenes from PEV, exerting a block to *cis* spreading of a chromatin state (24, 25). To date, a set of insulators have been identified in *Drosophila,* including BEAF-32 (Boundary element associated factor of 32 kDa), dCTCF (*Drosophila* homolog of CTCF), Su(Hw) (Suppressor of hairy wing), Zw5 (Zeste-white-5), GAF (GAGA factor) and recently described Pita and ZIPIC (zinc-finger protein interacting with CP190) (26, 27). Numerous studies demonstrated that insulators are responsible for a vast number of genomic functions, including stimulation of gene transcription, enhancer-blocking and barrier insulation partitioning of eukaryotic genomes into independently regulated domains (27–29). Hence, one may hypothesize that gene loci capable of adaptation to heterochromatin probably share specific sites of insulation that ensure their expression in the repressive environment.

To address this issue, we investigated molecular evolution of *Myb* and *Ranbp16* genes in the *Drosophilidae* lineage. *Myb* is an essential gene encoding a transcription factor involved in transition from G2 to M phase of the cell cycle (30, 31). *Ranbp16* encodes a RanGTP-binding protein belonging to the importin-β superfamily and mediates translocation of proteins into the nucleus. Both genes are located in euchromatic loci of the *D. melanogaster* X-chromosome, while in other studied *Drosophila* species belonging to Sophophora and Drosophila subgenus they are found in genomic regions with a high density of repetitive DNA elements upstream, downstream and within introns, suggesting their location in pericentric heterochromatin. Regardless of the euchromatic or heterochromatic surroundings, the promoter region of *Myb* displays high sequence homology and stable structural organization among *Drosophila* species studied so far. We find that the conserved motifs in the promoter sequence of *Myb* serve as a binding site for the chromatin insulator protein BEAF-32 and transcriptional factor DREF (The DNA replication-related element (DRE)/DNA replication-related element-binding factor). Furthermore, we demonstrate that BEAF-32 occupies not only the promoter region of *Myb* in two evolutionarily distant species, *D. melanogaster* and *D. virilis*, but also, in cooperation with other insulators dCTCF and GAF, is present in the promoters of most studied heterochromatin genes described in *D. melanogaster*. Importantly, the evolutionary gene repositioning between euchromatin and the pericentric heterochromatin occurred with preservation of sites of insulation, keeping the binding of BEAF-32 in close proximity to the transcription start sites of these genes. Overall, we propose that insulator proteins, in particular BEAF-32, contribute to expression of heterochromatic genes and make them predisposed for evolutionary repositioning into transcriptionally repressive genomic environments.

## Results

### Evolutionary repositioning of *Myb* and *Ranbp16* genes between euchromatin and heterochromatin in *Drosophila* phylogeny

In order to determine whether *Myb* and *Ranbp16* gene locations have been rearranged on the evolutionary timescale, we first mapped these genes onto genomic scaffolds of *Drosophila* species separated by evolutionary distances from 5 to 40 million years (32–36). These include species of the melanogaster group (*D. melanogaster* and *D. yakuba*) and the obscura group (*D. persimilis* and *D. miranda*), with both groups belonging to the Sophophora subgenus, along with the virilis group (*D. virilis* and *D. novamexicana*) and the repleta group (*D. mojavensis* and *D. hydei*) that belong to the Drosophila subgenus. Next, we performed comparative analysis of *Myb* and *Ranbp16* genes, as well as the intergenic regions between these genes. As indicated in Fig. 1, the coding sequences of *Myb* (red nodes) and *Ranbp16* (blue nodes) genes are highly homologous between the species studied (Fig. 1). However, the regions between *Myb* and *Ranbp16* genes differ significantly among *Drosophila* species studied, exhibiting sequence homology mostly within the related groups. For instance, while *Myb* and *Ranbp16* genes of *D. melanogaster* and *D. yakuba* are embedded within a large gene cluster, their orthologues in other *Drosophila* species reside in the genomic regions mostly occupied by repetitive sequences, including the introns of *Ranbp16* gene (Fig. 1). Thus, one may conclude that *Myb* and *Ranbp16* genes in the species belonging to the melanogaster group are located in euchromatin, in the region containing a large gene cluster. On the other hand, the localization of the studied genes in the environment packed with repetitive elements and apparently representing a pericentric heterochromatic region is typical for most other *Drosophila* species studied here including the virilis, obscura and repleta groups (Fig. 1). According to the integrated physical and cytogenetic map of Schaeffer et al. (37), as well as overall homology of the studied regions, *Myb* and *Ranbp16* genes are located at the chromosome X in all *Drosophila* species studied.

**Fig. 1.**
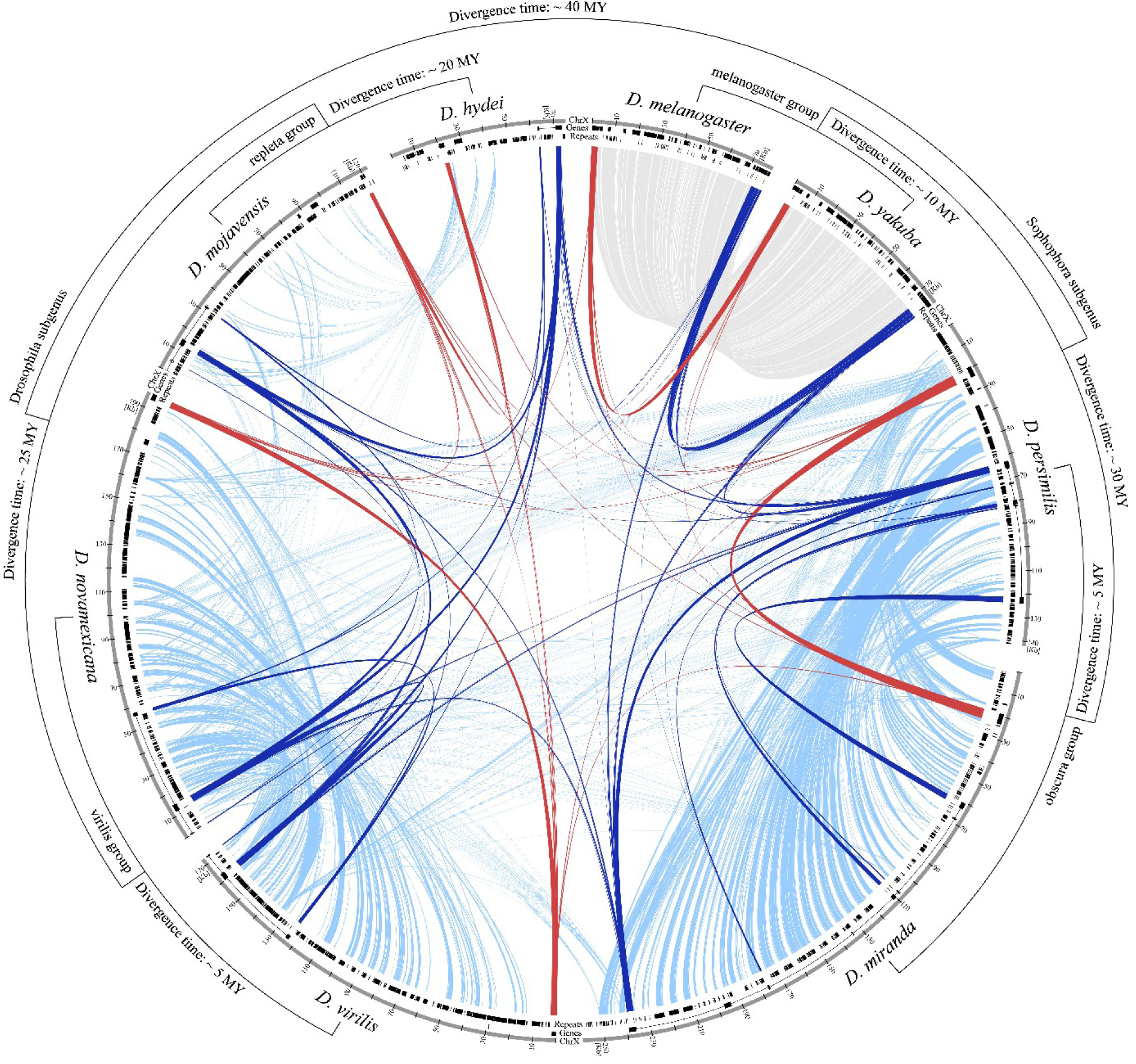
Circular plot demonstrating the overall homology of genomic regions comprising *Myb* and *Ranbp16* genes in the species of *Drosophila* genus. Color nodes indicate homology of *Myb* (red nodes) and *Ranbp16* (blue nodes) and intergenic regions between these genes consisting of repeats (light blue nodes) and protein-coding genes (gray nodes) among the species of the Sophophora and Drosophila subgenera. Tracks of the plot indicate the region of comparison on chromosome X, coordinates of genes including *Myb* and *Ranbp16,* as well as the content of repeats in the plotted region. The phylogenetic tree indicates the relationships among species with estimated time of divergence according to Clark et al. (32), to Gao et al. (33) for the obscura group, O’Grady et al. (36) for the virilis group and Gibbs et al. (34) for the repleta group.

Next, we studied in more detail the genomic loci containing *Myb* and *Ranbp16* genes, focusing on two evolutionarily distant species, *D. melanogaster* and *D. virilis,* separated by 40 million years of evolution (32). The single copies of *Myb* and *Ranbp16* genes map to the chromosome X of *D. melanogaster* at the cytogenetic loci 13F14 and 14A1 of salivary gland polytene chromosomes, respectively (Fig. 2A). These regions are significantly less enriched with heterochromatic marks H3K9me3 and HP1a than telomeric and pericentric regions of the chromosome and, hence, represent typical euchromatin. As mentioned earlier, *Myb* and *Ranbp16* genes in *D. melanogaster* are located at a distance ∼ 80 Kb from each other within a large protein-coding gene cluster, which includes only a few repetitive sequences (Fig. 2A).

**Fig. 2.**
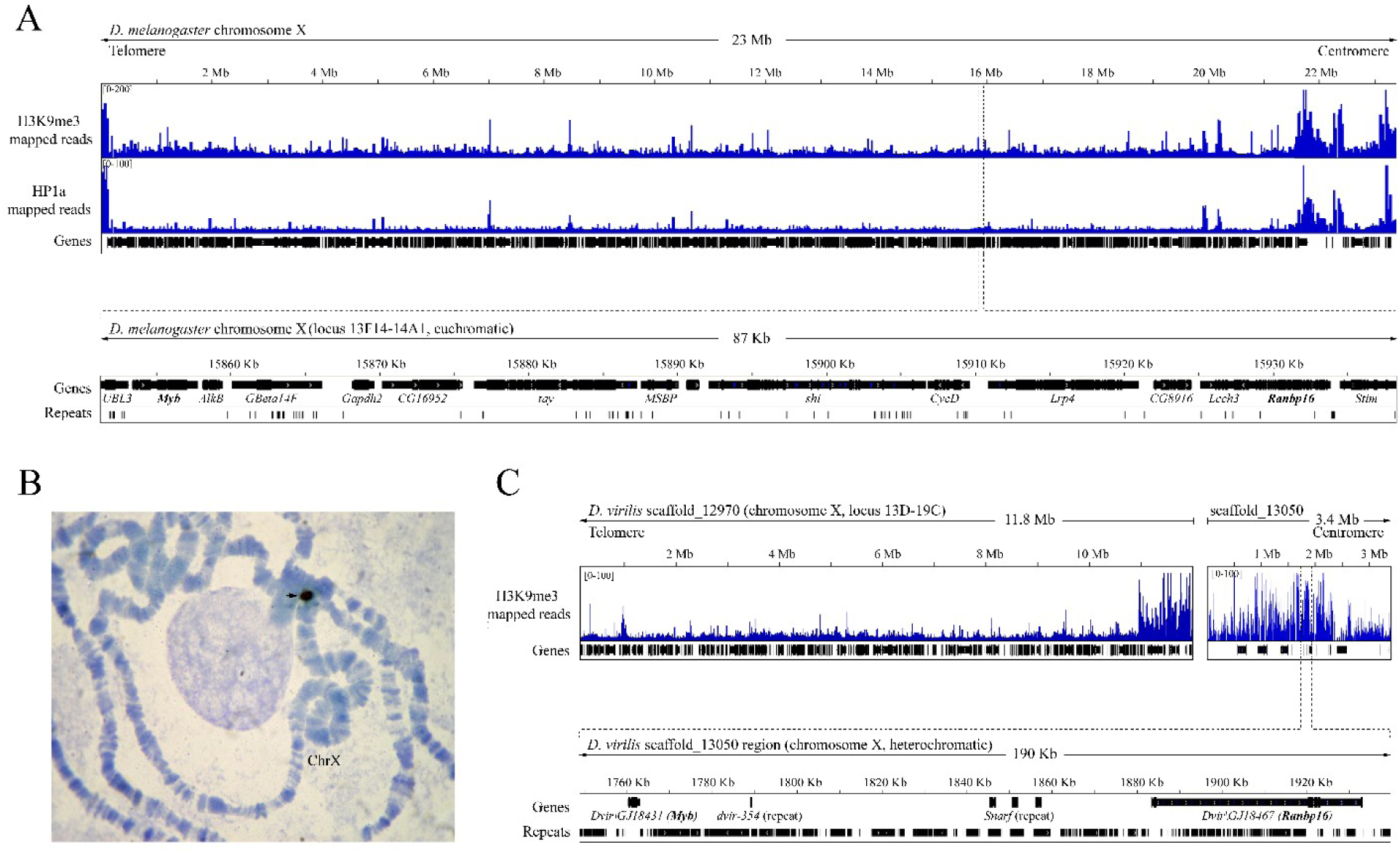
Analysis of genomic regions comprising *Myb* and *Ranbp16* genes in *D. melanogaster* and *D. virilis*. A) Genomic map of whole assembled chromosome X of *D. melanogaster* with mapped ChIP-seq reads of heterochromatic markers H3K9me3 and HP1a and the region depicting *Myb* and *Ranbp16* gene location (studied genes are marked with bold font). B) DNA *in situ* hybridization of *Myb* gene probe to polytene chromosomes of *D. virilis*. Black arrow indicates the hybridization signal at the base of chromosome X of *D. virilis*. C) Genomic map of scaffold_13050 of *D. virilis* and the proximal scaffold_12970 which is attributed to chromosome X of *D. virilis* according to Schaeffer et al. (37) with mapped ChIP-seq reads of H3K9me3 and the region containing *Myb* and *Ranbp16* genes (marked with bold font) on the larger scale.

To confirm that *Myb* and *Ranbp16* reside in a heterochromatic region in *D. virilis,* we performed DNA *in situ* hybridization on polytene chromosomes using the unique sequence of *Myb* gene as a probe. *In situ* hybridization indicates that *Myb* is located at the base of chromosome X near the chromocenter in *D. virilis* (Fig. 2B). Mapping analysis of *Myb* and *Ranbp16* genes on the genomic scaffolds of *D. virilis* reveals that both genes reside in one scaffold_13050 at a distance ∼ 120 Kb from each other (Fig. 2C). In contrast to *D. melanogaster*, the region between these genes in *D. virilis* does not contain any other protein-coding genes and consists of remnants of TEs and other repeats diverged to varying degrees (Fig. S1). We used annotation of known genomic scaffolds of *D. virilis* made by Schaeffer et al. (37) to assign the proximal scaffold of *D. virilis* genome r1.06 to the centromeric region of chromosome X. According to specific marker genes, we retrieved scaffold_12970 and extended it with scaffold_13050 containing *Myb* and *Ranbp16* genes, keeping some space unassembled between these scaffolds (Fig. 2C). Enrichment profile of H3K9me3 clearly indicates that the putative euchromatin-heterochromatin border lies in the proximal 1Mb of scaffold_12970 (Fig. 2C). The whole scaffold_13050, including the region where *Myb* and *Ranbp16* are localized, is heavily occupied by the H3K9me3 mark in comparison with the most contiguous fragment of scaffold_12970 (Fig. 2C).

Taken together, our data indicate that the essential *Myb* and *Ranbp16* genes reside in contrasting chromatin regions, euchromatic or heterochromatic, in the species of the *Drosophila* genus studied so far.

### Both *Myb* and *Ranb16* genes are under purifying selection regardless of their heterochromatic or euchromatic localization

Next, we examined whether coding sequences of *Myb* and *Ranbp16* underwent negative (purifying) or positive selection during evolution and what was the impact of heterochromatic location on molecular evolution of these genes. To this end, we first estimated the number of base substitutions per site in their coding sequences (Fig. S2A, B), and then calculated the ratio of non-synonymous to synonymous substitutions (dN/dS) (Fig. S2C, D) using eight aforementioned representatives of the Sophophora and Drosophila subgenera, as well as the sequences from *Anopheles gambiae* as an outgroup. The results suggest that both *Myb* and *Ranbp16* genes are under purifying (negative) selection (dN/dS < 0.2 in all pairs of comparison) when present either in heterochromatin or euchromatin (Fig. S2).

We also performed multiple alignment of protein sequences of Myb and Ranbp16 from Diptera species and their orthologs in mouse and human. The results show a high degree of conservation of both proteins even between dipteran and mammalian species (Fig. S3). The consensus sequences of aligned proteins are properly recognized as Helix-Turn-Helix (HTH) DNA-binding domain and Importin-β domain that characterize the described structure of Myb and Ranbp16 proteins, respectively (Fig. S3).

### Evolutionary changes in gene organization of *Myb* and *Ranbp16* in *Drosophila* species

To study how the structure of *Myb* and *Ranbp16* genes has been changed in the course of evolution, resulting in their placement in different chromatin contexts, we compared the structure of these genes in eight *Drosophila* species.

As a starting point, we retrieved the available sequences of *Myb* and *Ranbp16* genes from *D. melanogaster*, *D. yakuba*, *D. persimilis*, *D. obscura*, *D. virilis*, *D. mojavensis* and *D. hydei* from Flybase and NCBI databases. To date, full annotation of genes including 5’ and 3’UTR (untranslated region) of most *Drosophila* species, except for *D. melanogaster,* is absent. To resolve this issue and obtain the most complete sequence of studied genes, we used sequences of closely related species (for example, *D. obscura* sequences were used to resolve the *D. persimilis* gene structure, etc.). Only those alignments that mapped on the same DNA strand adjacent to the existing annotated sequence with E-value > e-80 were considered as relevant and used for reconstruction of gene structure. Also, 5’RACE (Rapid amplification of cDNA ends) was performed to determine the missing 5’UTR of *Ranbp16* gene of *D. virilis*. Due to the lack of annotation, the sequences of *Myb* and *Ranbp16* genes of *D. novamexicana* were reconstructed using the sequences of related *D. virilis* species and blastn algorithm. Gaps in the continuous alignment were considered as introns. All information, including accession numbers and genomic coordinates of the studied genes in *Drosophila* species, is listed in supplementary table S1.

Despite prolonged evolution, the structure of *Myb* gene was preserved virtually unchanged in terms of overall gene length and exon-intron organization (Fig. 3A). A single intron located in the protein-coding region of this gene is conserved both in position and approximate size in the species of both subgenera (Fig. 3A). However, sequence homology of this intron is conserved only within species belonging to one subgenus, Sophophora or Drosophila. This suggests that the intronic sequence *per se* probably does not affect gene regulation. Therefore, the heterochromatic *Myb* gene tends to be stable during evolution of the Drosophila genus and retains the single intron of approximately the same size in different species.

**Fig. 3.**
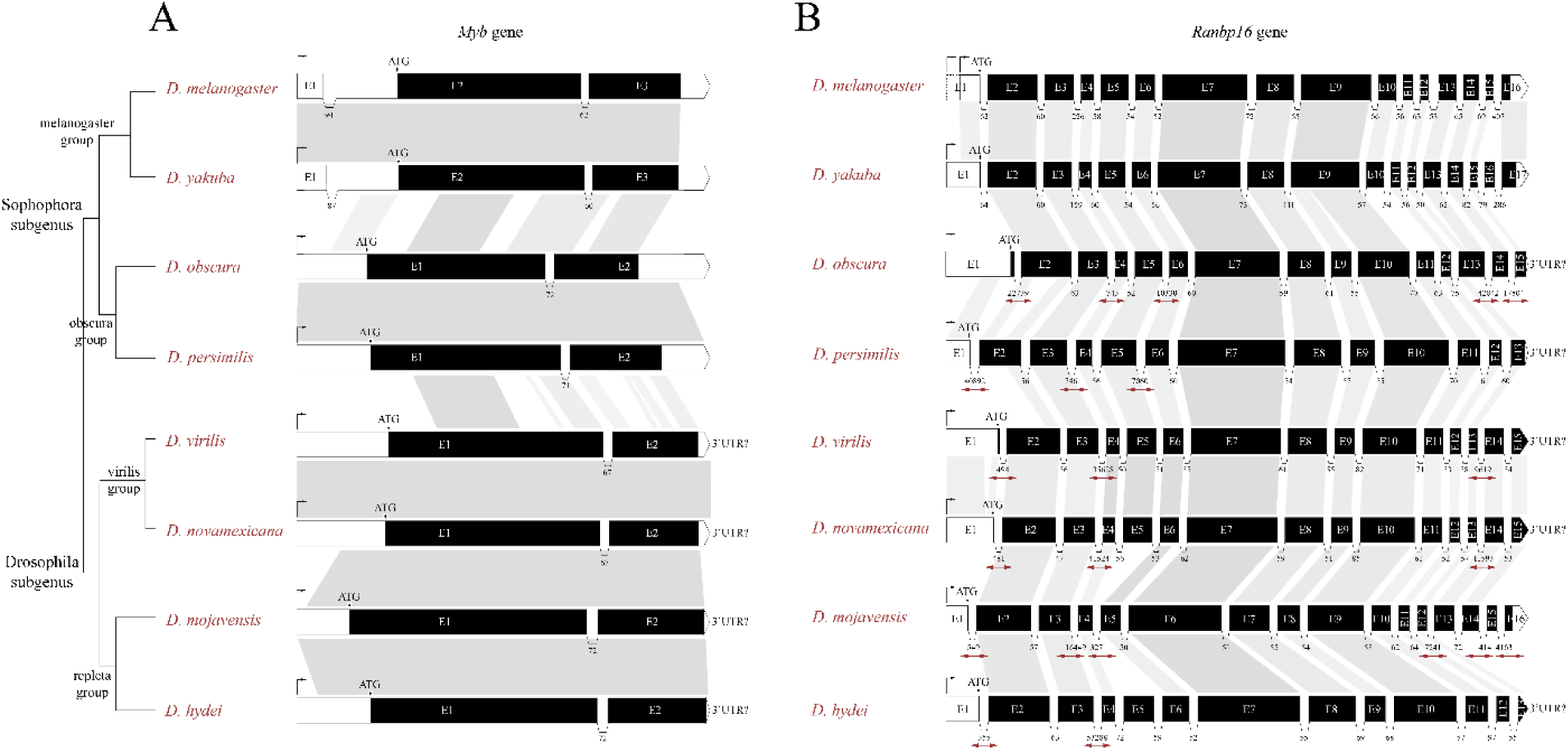
Comparative gene organization of *Myb* (A) and *Ranbp16* (B) genes among eight *Drosophila* species. White and black regions on the scheme denote untranslated and coding regions, respectively. The gray range between genes indicates homologous regions. Homology of introns is showed only for the *Myb* gene. Rightwards corner arrow at the beginning of genes indicates the transcriptional start site. Numbers beneath the introns reflect their size in base pairs. Introns that have been increased in size by transposable element insertion are highlighted by red bidirectional arrows. The 3’untranslated regions (3’UTR) are not given for most of the genes due to the absence of their annotations and homology to closely related species. Bootstrap consensus phylogenetic tree is given according to Clark et al. (32), Gao et al. (33), O’Grady et al. (36) and Gibbs et al. (34) considering different groups of the *Drosophila* genus.

In contrast to *Myb*, the structure of *Ranbp16* is more complex and demonstrates greater flexibility in terms of extension of total gene length by increasing the size of its introns (Fig. 3B). The length of *Ranbp16* gene in the species of melanogaster group (*D. melanogaster* and *D. yakuba*) is ∼ 6700 bp and includes up to 15 introns not exceeding 407 bp in length. In the course of evolution, the gene structure of *Ranbp16* in other studied *Drosophila* species was expanded to varying degrees by multiple insertions of TEs. For example, the length of *Ranbp16* gene is approximately 100 Kb in *D. obscura*, 55 Kb in *D. virilis* and 40 Kb in *D. mojavensis* (Fig. 3B). Interestingly, the introns prone to TE insertions increasing the length of the gene are strictly defined. They include the first, third and fifth introns from the TSS (transcriptional start site) of the gene, as well as a few introns at the distal end of the gene (Fig. 3B). The extension of introns located in the middle of the gene was not observed in any of the studied species, suggesting that these introns may have a structural or regulatory role in *Ranbp16* function.

Thus, the structure of heterochromatic genes may exhibit great flexibility in terms of overall gene length extension, if this extension does not disrupt gene activity.

### Promoter of *Myb* gene contains motifs for binding of chromatin remodeling proteins in *Drosophila*

To reveal specific properties of gene loci allowing the essential *Myb* and *Ranbp16* genes to be actively transcribed in both euchromatin and heterochromatin, we studied the promoter region of *Myb* by searching for common and different motifs in *Drosophila* species. For this purpose, we expanded the list of analyzed species to 19 representatives of the Sophophora and 11 representatives of the Drosophila subgenera. As mentioned above, due to the absence of gene annotation for a range of studied species (*D. miranda*, *D. guanche*, *D. subobscura* and all virilis group species with the exception of *D. virilis*) the putative TSS was set as the first mapped nucleotide of 5’UTR of *Myb* gene of related species. Due to the insertion of DAIBAM MITE element at the position 92 bp upstream of the TSS of *Myb* gene in *D. virilis*, the analyzed region was shortened to 100 nt for all species. Using the MEME suite (38) we were able to identify three motifs that are present in the promoter of *Myb* gene in all analyzed species of *Drosophila* (Fig. 4A). Search through OnTheFly (39) and REDfly v5.6 (40, 41) databases of known transcription factors and their binding sites indicates that the two highest-scoring motifs contain a potential binding site for insulator protein BEAF-32 and transcriptional factor DREF (Fig. 4B). The third motif shows limited homology to the binding site of ecdysone receptor co-activator Taiman and another undescribed factor (Fig. 4B).

**Fig. 4.**
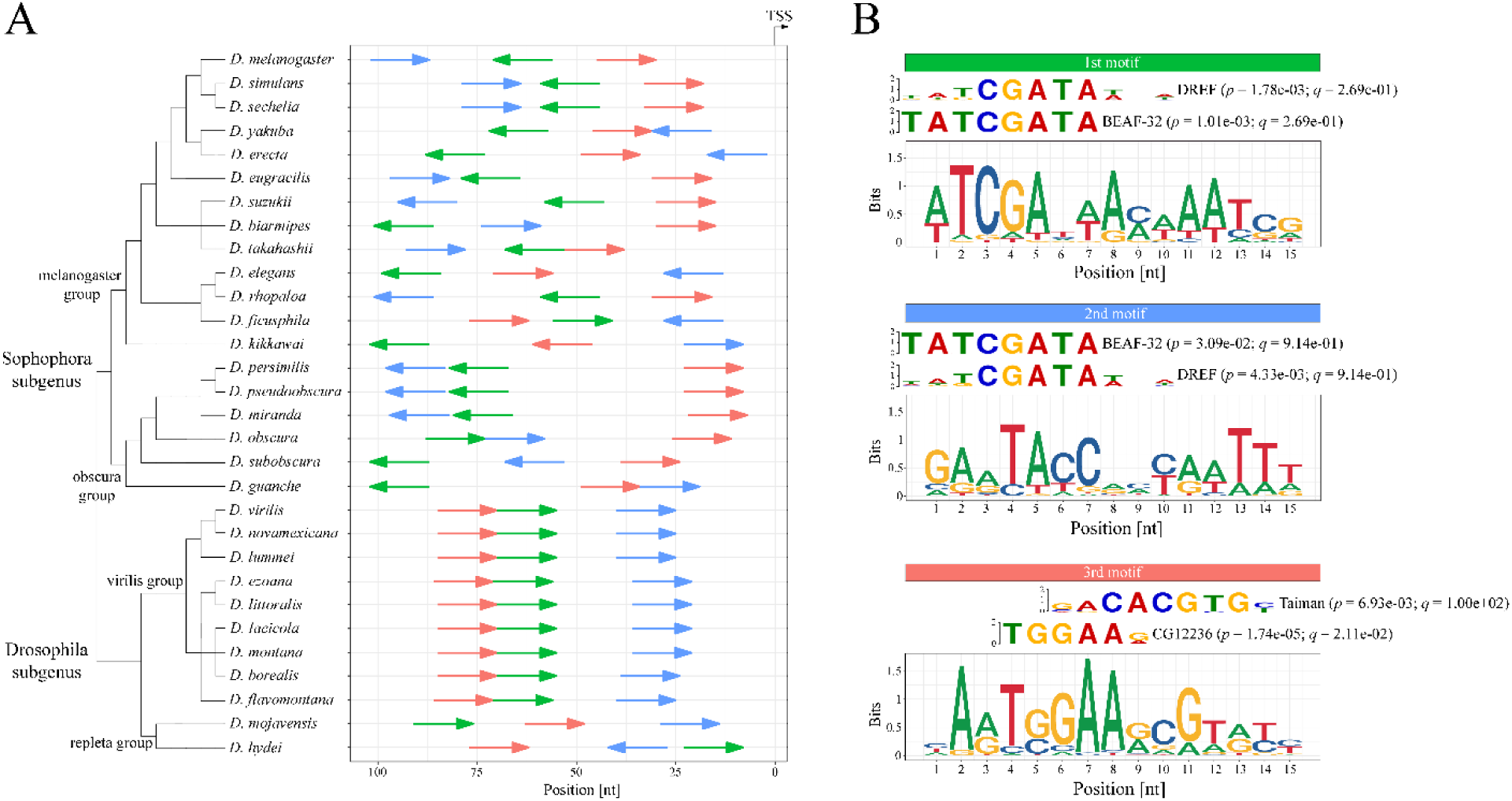
Comparative promoter analysis of *Myb* gene in 30 *Drosophila* species. A) Distribution of common motifs and their orientation (shown by colored arrows) in the promoter of *Myb*. Prior to the analysis, orientation of all sequences was adjusted so that the transcription start site (TSS) would be on the right. B) Sequence logos of the common motifs among the studied *Drosophila* species with matches between motifs and binding sites of known transcription factors. Logos for binding sites of the factors are shown with the corresponding *p*- and *q*-values. The colors of motifs in A) and B) correspond to each other. Bootstrap consensus phylogenetic tree is given according to Clark et al. (32), Gao et al. (33), O’Grady et al. (36) and Jezovit et al. (35).

### Insulator protein BEAF-32 is enriched in promoters of *Myb* and *Ranbp16* genes in both *D. melanogaster* and *D. virilis* species

To demonstrate that the insulator protein BEAF-32 binds to the promoter region of *Myb* gene, we used ChIP-seq data to profile BEAF-32 occupancy in the gene loci containing *Myb* and *Ranbp16* genes in *D. melanogaster* and *D. virilis*. Furthermore, we applied ChIP-seq to profile RNA polymerase II (Pol II) distribution in the promoters of *Myb* and *Ranbp16* genes. Additionally, we used ATAC-seq data to correlate BEAF-32 and Pol II enrichment with the nucleosome-free conformation of chromatin in the promoters of these genes indicative of intensive transcription. Mapping of RNA-seq and H3K9me3 ChIP-seq data also provides valuable information regarding expression levels of these genes and the heterochromatic profile of the analyzed gene loci, respectively.

As seen in Fig. 5A, in *D. melanogaster* the locus containing *Myb* gene and adjacent *AlkB* gene is highly enriched with BEAF-32, with the peak summit of mapped BEAF-32 reads in the promoter of the *Myb* gene (Fig. 5A). The enrichment of Pol II and mapped ATAC-seq reads reside within the region starting from the peak of BEAF-32 binding to the TSS of *Myb,* indicating that BEAF-32 binds to DNA at the boundary of the *Myb* gene locus (Fig. 5A). Likewise, the promoter of *Ranbp16* gene of *D. melanogaster* is enriched with BEAF-32, located slightly upstream from the Pol II binding site and open chromatin region (Fig. 5B).

**Fig. 5.**
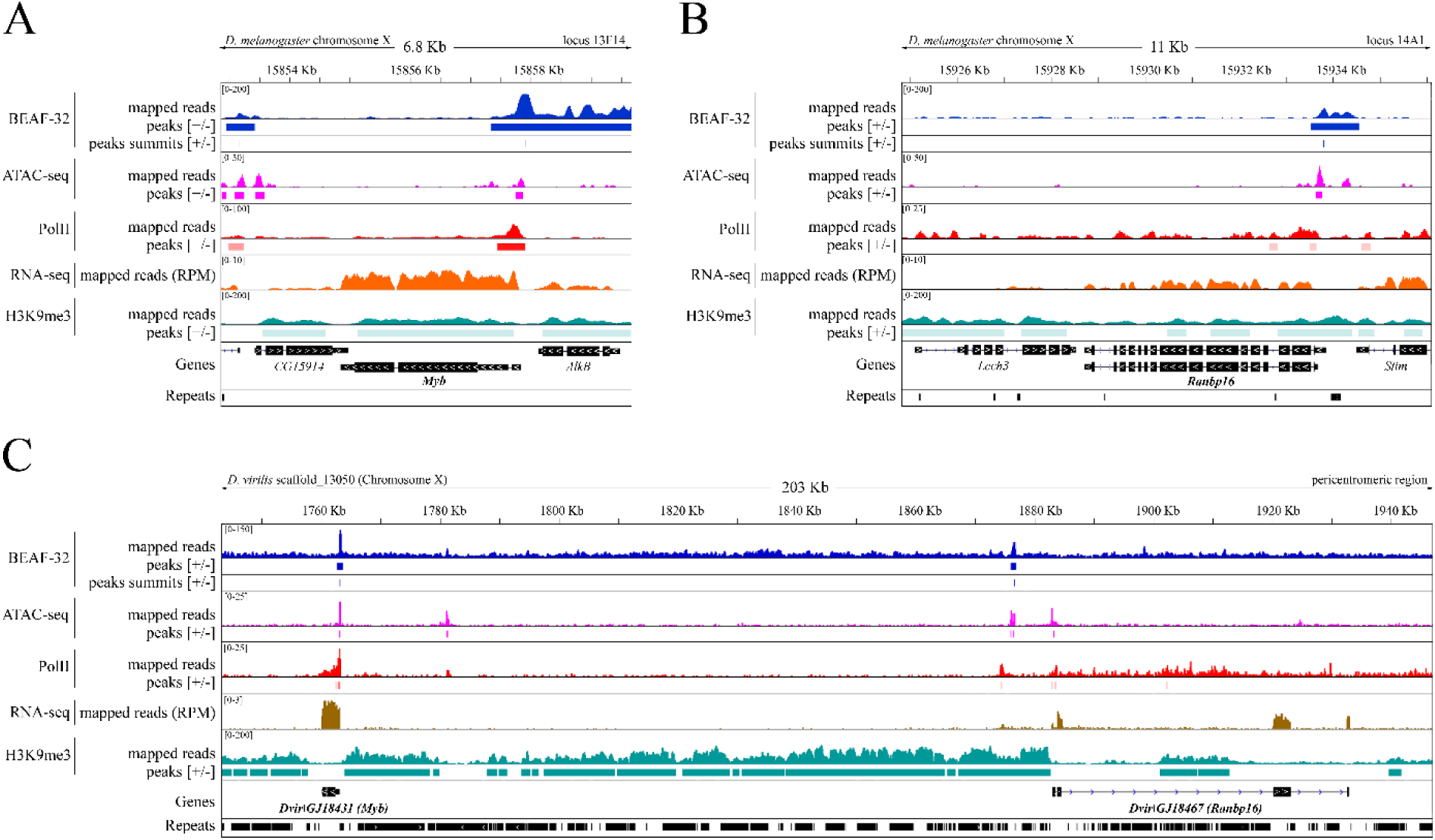
Enrichment profiles of BEAF-32, RNA Pol II, H3K9me3, RNA-seq and ATAC-seq reads in the genomic regions comprising *Myb* and *Ranbp16* genes in *D. melanogaster* and *D. virilis*. A) and B) Enrichment of ChiP-seq reads within *Myb* and *Ranbp16* genic loci in *D. melanogaster*. C) The enrichment profile within the *Myb* and *Ranpb16* genic loci and the intergenic regions between these genes in *D. virilis*. Mapped ChIP-seq reads without normalization (mapped reads), calculated areas of enrichment relative to the input data (peaks; *p*- and *q*-value < 0.05) are shown for BEAF-32, Pol II, H3K9me3 and ATAC-seq data. Summits of the enriched reads are shown for BEAF-32 (peaks summits). RNA-seq reads were normalized to the sequence depth (RPM, reads per million).

A similar pattern of BEAF-32 occupancy in the promoters of heterochromatic *Myb* and *Ranbp16* genes is observed in *D. virilis* (Fig. 5C). However, the enrichment profile of BEAF-32 upstream of the TSS of *Ranbp16* gene in *D. virilis* and *D. melanogaster* has one difference which is worth mentioning. In contrast to *D. melanogaster*, where BEAF-32 is enriched in close proximity to the TSS of *Ranbp16*, the binding of BEAF-32 to DNA in *D. virilis* is observed only at a distance of ∼ 6.5 Kb from the TSS of *Ranbp16* (Fig. 5). Within this range, a couple of transposon insertions are located. According to the data obtained by CAGE-seq (Cap Analysis Gene Expression), *Ranbp16* gene in *D. melanogaster* has two TSS located at a distance of ∼ 200 bp from each other, giving rise to slightly distinct transcripts in terms of the length of their 5’UTR. Transcripts with the shorter 5’UTR are transcribed throughout the development of *Drosophila*, whereas transcripts with the longer 5’UTR are present mostly at the larval stage (Fig. S4). Given that BEAF-32 is enriched within the upstream promoter of *Ranbp16* in *D. melanogaster*, we assumed that in *D. virilis* the distance to the upstream promoter has been extended due to transposon insertions that may be spliced in the course of transcription. However, we failed to observe more than a single TSS by 5’RACE analysis at the larval and imago stages as well as in the gonads of *D. virilis,* suggesting either loss of the second promoter or its extremely low expression level.

Overall, these results indicate that the insulator protein BEAF-32 is enriched in the vicinity of the promoters of *Myb* and *Ranbp16* genes embedded in distinct chromatin structures (euchromatic vs heterochromatic) in *D. melanogaster* and *D. virilis*, respectively.

### Binding of BEAF-32 in the proximity of the transcription start sites is preserved in the course of evolutionary repositioning of gene loci between euchromatin and heterochromatin

Considering the above results, two important questions may be asked. First, do the BEAF-32 binding sites in the proximity of promoter regions represent a peculiar property of *Myb* and *Ranbp16* genes, or are they a common feature of heterochromatic genes? Second, do the BEAF-32 binding sites found in these promoters emerge in the course of adaptive evolution of genes transposed to heterochromatin? Alternatively, they might represent an ancestral feature which favors repositioning to the repressive environment without any deleterious impact on fitness.

To address these questions, we analyzed a representative set of more than 30 genes that reside in the pericentric heterochromatin of both arms of chromosome 2 in *D. melanogaster,* while in *D. virilis* the orthologs of these genes are clustered in different loci located in the euchromatin of the same chromosomal elements (Table S2) (21). Genes demonstrating the opposite scenario of repositioning were also included in the analysis. Among them we considered the heterochromatic genes *Stim* and *Rrp47* that are located on the same scaffold_13050 as *Myb* and *Ranbp16* of *D. virilis,* as well as the genes *RpL15*, *Calr*, *Atg2* and *CG40228* that are embedded in scaffold_12736 located near the chromocenter of *D. virilis* (Table S2) (18). In contrast to *D. virilis*, most of these genes with the exception of *RpL15* and *CG40228* are located in euchromatic regions of different chromosomal elements in *D. melanogaster* (Table S2).

Using these sets of genes, we performed enrichment analysis of BEAF-32, Pol II, H3K9me3 and ATAC-seq reads upstream and downstream of TSS of all these genes (Fig. 6). As indicated in Fig. 6A, the insulator protein BEAF-32 is enriched in the proximity of TSS or within 10 Kb upstream of TSS of both heterochromatic (Group 1) and euchromatic genes (Group 2) of *D. melanogaster* (Fig. 6A). The binding area of BEAF-32 is strongly correlated with the enrichment profile of Pol II and ATAC-seq reads (Fig. 6A). Importantly, analysis of *D. virilis* genes revealed a predominantly similar enrichment profile of BEAF-32 in both clusters of genes partitioned with regard to their heterochromatic or euchromatic location in the genome (Fig. 6B).

**Fig. 6.**
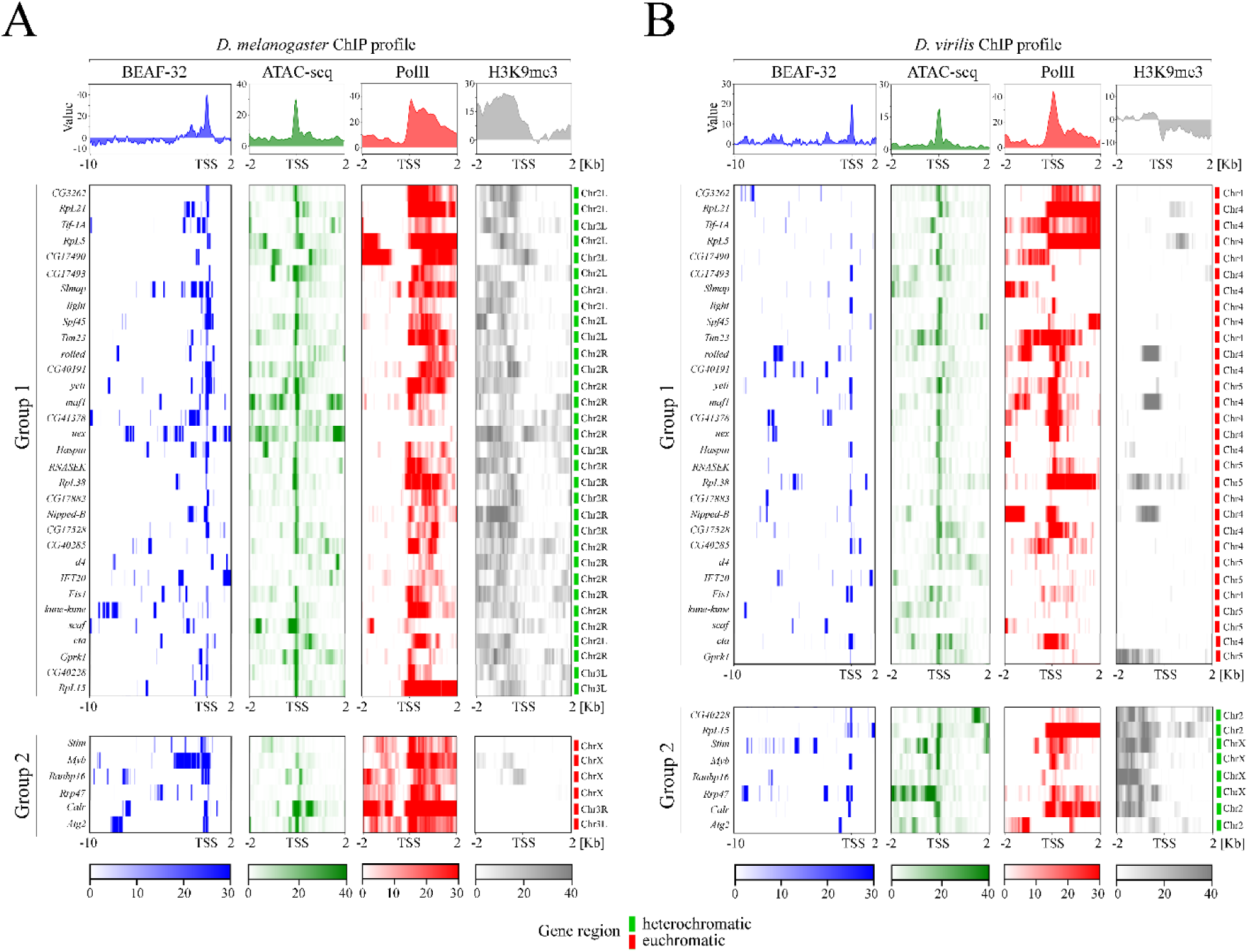
Enrichment profiles and heatmaps of BEAF-32, RNA Pol II, H3K9me3 and ATAC-seq reads upstream and downstream of the transcriptional start site (TSS) of genes that were shown to be repositioned from euchromatin to heterochromatin during *Drosophila* evolution. A) and B) show the enrichment profiles of ChIP-seq reads of genic loci of *D. melanogaster* and *D. virilis*, respectively. Gene loci are subdivided into groups 1 and 2 corresponding to their heterochromatic (marked by green color) and euchromatic (red color) location. Note that the analyzed area for BEAF-32 enrichment includes 10 Kb upstream and 2 Kb downstream from TSS, while for RNA Pol II, H3K9me3 and ATAC-seq reads area includes 2 Kb both upstream and downstream from the TSS. Color codes below the heatmaps indicate fold enrichment.

Taken together, these results indicate that BEAF-32 binding sites are present in the proximity of the promoters of genes that were repositioned into heterochromatin during the evolution of the *Drosophila* genus.

### Boundaries of heterochromatic gene loci are occupied by diverse insulator proteins in *D. melanogaster*

As demonstrated above, the insulator protein BEAF-32 is highly enriched in the proximity of promoters of heterochromatic genes. To further elucidate how widespread are insulators in heterochromatic regions, we analyzed the enrichment profiles of other *Drosophila* insulators, such as dCTCF, GAF, Pita, ZIPIC, Su(Hw) and Zw5, in the pericentric region of chromosome 2L of *D. melanogaster*. We find that, similar to BEAF-32, dCTCF and GAF have multiple binding sites, while Pita, ZIPIC and Su(Hw) are only modestly represented in the pericentric heterochromatin (Fig. S5A). Finally, Zw5 binding sites are completely absent from the studied region (Fig. S5A).

Next, we analyzed the enrichment profile of insulators upstream, downstream and within the bodies of heterochromatic genes located in the pericentric region of *D. melanogaster* chromosome 2L (Fig. S5B). We observed that BEAF-32 is not the only insulator found in the vicinity of TSS in heterochromatic genes. Although to a lesser extent, the insulators dCTCF and GAF are also enriched at the TSS of these genes (Fig. S5B). Interestingly, a few binding sites for Pita are present at the boundaries of gene loci as indicated by binding near the transcriptional start and transcriptional end sites (TES). Despite the binding of ZIPIC and Su(HW) to the pericentric DNA, the sites of their binding are not localized at the TSS and TES of the genes, suggesting that these insulators may be involved in long-range interactions of chromosomes, and hardly have any direct impact on gene expression (Fig. S5B).

Overall, these data suggest that the insulator proteins are involved in the maintenance of pericentric heterochromatin conformation and normal functioning of structural genes located in pericentromeric heterochromatin.

## Discussion

During speciation, the genomes of *Drosophila* species underwent multiple chromosome rearrangements that disrupt gene order, modifying or preserving gene function (17, 42). In this study, we show that in the course of evolution of *Drosophila* species the essential *Myb* and *Ranbp16* genes have been repositioned into different chromatin types, i.e. euchromatin or heterochromatin. Despite the contrasting chromatin structure and local repressive environment of heterochromatic regions enriched with repetitive DNA, both genes were shown to be under purifying selection due to their highly conserved function. However, the overall gene organization may not be as conserved as the protein-coding region of heterochromatic genes. It was reported in a number of studies (18–21), that genes located in heterochromatic environment may increase their size by becoming targets for multiple TE insertions in their introns. Similar to the described evolutionary history of *RpL15* and *Yeti* (18, 19), the organization of *Myb* remained stable during evolution and includes a single short intron in the coding region of all species belonging to Sophophora and Drosophila subgenera. In contrast to *Myb*, the structure of *Ranbp16* gene embedded in heterochromatin has evolved to accumulate numerous TEs in its introns. These introns extended the gene length from ∼40 kb in *D. mojavensis* to ∼180 kb in *D. miranda*. The adaptive value of different intron lengths in the same gene in related species is unknown. A simple explanation of different intron content of the studied genes (*Myb* vs *Ranbp16*) stems from the fact that selective pressure favors short introns in highly expressed genes rather than in genes that are expressed at lower levels (43). In addition, a phenomenon called intron delay may limit the ability of cells with a short mitotic cycle to transcribe long primary transcripts (44, 45). On this view, *Myb,* which is a highly expressed gene throughout *Drosophila* development, retained its organization regardless of its euchromatic or heterochromatic location. On the contrary, *Ranbp16* gene, one of the members of importin-β family, is expressed at low level and probably exhibits tissue-specific patterns of expression. This may result in its more labile organization on the evolutionary timescale, with a remarkable increase in size in the heterochromatin context.

An unexplained question is the differential susceptibility of *Ranbp16* introns to expansion. A dramatic increase of *Ranbp16* gene length in several species occurred only due to the extension of certain but not all introns comprising the gene. It may be thought that selective pressure affects different parts of *Ranbp16* gene to varying degrees, enabling expansion of some introns and preserving the length of others. In genomes of animals, plants, fungi, and protists, intron positions could be conserved throughout prolonged evolutionary times, suggesting their involvement in gene expression and regulation, including mRNA processing (46–48). Numerous studies in animals and plants have emphasized that introns may carry functional *cis*-acting elements affecting gene expression even in the absence of promoter sequences (49–53). Recently, it was shown that nucleosomes preferentially occupy exons rather than introns, and this phenomenon seems to be interconnected with specific histone modifications favoring gene expression (54–56). Thus, conservation of specific introns in the evolution of heterochromatic genes suggests their role in regulation of expression of these genes. However, the cross-talk between regulatory elements in introns, gene architecture, chromatin structure, and nucleosome positioning and modifications, especially in the heterochromatin, needs to be investigated in more detail.

According to studies of PEV, genes that are juxtaposed with heterochromatin by chromosomal rearrangements or transposition events exhibit a variegating phenotype resulting in silencing of genes due to heterochromatin environment (8). Given the peculiarities of heterochromatic genes, such as accumulation of TEs within their introns, one may suggest that heterochromatic genes have evolved to utilize the advantage of the heterochromatic environment and became dependent on heterochromatin specific proteins, such as HP1a (4, 57–60). However, as was demonstrated previously, the regulatory regions including gene promoters apparently did not undergo drastic changes during their evolution in the heterochromatic environment (20). Moreover, evolutionary repositioning preferentially occurred only with gene clusters exhibiting close association with HP1a, suggesting that HP1a binding to these genes was present in the progenitor (21). Taken together, these observations allow one to suggest that certain gene loci are favored for repositioning from euchromatin to heterochromatin in the course of evolution. If this is the case, gene loci relocated to heterochromatin probably should retain the transcriptionally active euchromatin-like structure of chromatin capable of efficient transcription in the heterochromatin. Indeed, the proximal regulatory regions of heterochromatic genes are not occupied by the heterochromatic mark H3K9me3, forming a nucleosome-free binding platform for transcriptional factors and RNA polymerase II (Fig. 5 and 6).

How might these observations be explained? What determines the “immunity” of regulatory regions of heterochromatic genes to the repressive surroundings? In this study, we have examined the promoter of *Myb* gene in thirty *Drosophila* species and revealed common motifs between these sequences that serve as binding sites for the insulator protein BEAF-32 and transcriptional factor DREF. Originally, insulators were defined as regulators of interaction between enhancers and promoters able to block heterochromatin spreading and PEV (24, 25). DREF is known to mediate activation of transcription of genes involved in DNA replication and cell proliferation, including *Myb* gene (61, 62). Furthermore, *cis*-acting elements that exercise the transcriptional control of genes by DREF are conserved between such evolutionarily distant species as *D. melanogaster* and *D. virilis* (*63*). The DNA recognition motif for DREF (CGATA) is similar to that of BEAF-32 (TATCGATA), and the binding of DREF to DNA has been shown to antagonize the binding of BEAF-32 *in vitro* (64). Recent studies of the relationship between BEAF-32 and DREF showed that DREF co-localizes at the same genomic sites as BEAF-32 and other insulator proteins and is enriched at the boundaries of topologically associated domains (TAD) (65). Together, these data suggest that DREF function is mediated by and probably depends on insulators on an evolutionary timescale.

A growing body of evidence suggests that insulators exercise diverse roles, including barrier function, and mediate short and long-distance chromosomal contacts at the genome-wide scale (66–70). Our enrichment analysis of ChIP-seq data indicated that the insulator protein BEAF-32 is enriched upstream of the TSS of heterochromatic genes in *D. melanogaster* and *D. virilis,* demarcating the euchromatin-heterochromatin border between the promoter and the surrounding heterochromatin (Fig. 6). Furthermore, using the same set of genes that reside in different types of chromatin in *D. melanogaster* and *D. virilis,* we show that BEAF-32 binding is preserved in the proximity of the promoter regions of these genes during evolution of different *Drosophila* species, suggesting that BEAF-32 binding is an ancestral property of these genes.

Along with BEAF-32, insulators dCTCF and GAF are also enriched at the TSS of heterochromatic genes, while the binding peaks for insulators Pita, ZIPIC and Su(Hw) are distributed across the pericentric region to a much lesser extent. The insulators dCTCF and GAF have many overlapping binding sites with BEAF-32 and DREF in the *Drosophila* genome (65). Furthermore, sites containing DREF without BEAF-32 are instead enriched with the Su(Hw) recognition sequences (65). Pita also belongs to a class of insulator proteins that preferentially bind to promoters near the TSS (71). ZIPIC is described as facilitator of long-distance chromosomal interactions (26, 71).

How this complex machinery works to facilitate continuous gene expression and chromosomal architecture remains an unresolved question. It is of note that a direct impact of insulator presence on gene expression has been established for the *D. melanogaster* GAGA factor (GAF) that resides in the *hsp70* promoter. GAF mediates the recruitment of chromatin remodeling factors, including SWI/SNF, the CHD, and the ISWI family complexes, that ensure formation of nucleosome-free region in the *hsp70* promoter (72–74). Studies of BEAF-32 demonstrated that most BEAF-associated genes are transcriptionally active or highly expressed and are associated with negative elongation factor NELF that stimulates transcription levels by inhibiting promoter-proximal nucleosome assembly (67, 75). This provides evidence that BEAF-32 facilitates high levels of gene expression.

Previously, it was shown that heterochromatic genes take advantage of repetitive surrounding and heterochromatin factors, such as HP1a, to facilitate their expression and probably long-distance interactions between enhancers and promoters (11, 22, 23, 76). On the other hand, an analysis conducted by Yasuhara et al (20) revealed that promoters of heterochromatic genes have not undergone major alterations after repositioning into the repetitive environment of heterochromatin rejecting the existence of heterochromatin-specific promoter. Along these lines, the observed binding of BEAF-32 in the proximity of gene promoters which underwent repositioning between euchromatin and heterochromatin in evolutionarily distant species of *Drosophila* is very intriguing. In cooperation with HP1a, the presence of BEAF-32 at gene promoters probably contributed to the formation of “proto-heterochromatic” gene loci in the ancestral species of *Drosophila* and thus facilitated their normal functioning in the heterochromatic environment. Recently, a remarkable peculiarity of HP1a binding at several gene loci has been described, whereby HP1a can be recruited to gene promoters independently of H3K9 methylation (77). Hence, one may hypothesize that BEAF-32 mediates the barrier function blocking the spreading of heterochromatin to the promoter regions of heterochromatic genes, while HP1a maintains a proper chromatin structure at and around such gene loci.

While it is clear that further studies are needed to elucidate all the factors required for normal gene functioning in the heterochromatic surroundings, our results provide an important insight into molecular mechanisms operating in the evolution of heterochromatic genes to provide their expression in the repressive environment.

## Materials and methods

### Drosophila genomes and sequence analyses

Drosophila genomes and gene sequences for comparative analysis were extracted from FlyBase and NCBI databases. Sequences of genomic regions containing *Myb* gene of virilis group species e.g. *D. lacicola, D. littoralis, D. borealis, D. flavomontana, D. lummei, D. ezoana* were fetched from unpublished data of Dr. Venera Tyukmaeva & Prof. Michael Ritchie from the University of St. Andrews, UK (personal communication). In the case of absence of gene annotation (e.g. for *D. guanche*, *D. subobscura*, most of the virilis group species), orthologs were retrieved with TblastN (78) using protein sequence of the most closely related species (i.e. *D. obscura* for *D. subobscura* and *D. guanche*, *D. virilis* for *D. lacicola* and other virilis species) as queries. All the query subjects mapped on the same DNA strand adjacent to each other with E-value > e-80 were considered as true and used for reconstruction of coding sequences. Sequences between mapped subjects were considered as introns. Putative transcriptional start sites (TSS) of poorly annotated genes were identified with blastN (79) using 1^st^ exon sequence of related species. Blast results with E-value > e-60 and adjacent to annotated coding sequence at a distance not exceeding 600 nt were considered as true and the 1^st^ mapped nucleotide as TSS. All essential information, including genes IDs, genomes IDs, and genomic coordinates of *Myb* and *Ranbp16* in all studied species, is listed in table S1. Orthologous sequences of *Myb* and *Ranbp16* genes of *Anopheles gambiae* were extracted from VectorBase (https://www.vectorbase.org/) by the numbers AGAP008160 – *Myb* and AGAP004535 – *Ranbp16*. Protein sequences of *Myb* and *Ranbp16* (also known as *Xpo7*) of mouse and human were extracted from UniProt (https://www.uniprot.org/). Protein motifs were scanned using the PROSITE database and methodology (80, 81).

Estimation of repeat content in intergenic regions and within studied genes was performed using RepeatMasker (82) and computationally predicted libraries of TEs generated with REaS (83) that are available in FlyBase (ftp://ftp.flybase.net/genomes/aaa/transposable_elements/ReAS/v2/consensus_fasta/). For repeat masking of *D. miranda* genome, we used consensus sequences of TEs of *D. pseudoobscura* and *D. persimilis*, and for *D. hydei* we applied the library of *D. mojavensis*. TEs were classified using RepeatClassifier implemented in RepeatModeler software (84).

Multiple sequence alignment was performed with ClustalW (85) and Clustal Omega (86) programs (https://www.ebi.ac.uk/Tools/msa/) for nucleotide and amino acid alignments, respectively. Multiple protein alignments were visualized with Jalview (87).

Exon-intron structure of studied genes were visualized using WormWeb exon-intron graphic maker (http://wormweb.org/exonintron). Visualization of blast results was done with Kablammo (88).

Circular plot was made using Circos visualization tool (89). Flanking regions of *Myb* and *Ranbp16* (20 Kb upstream and downstream from the gene location) were used instead of intergenic regions, due to the long distance between these genes in *D. miranda* (> 17 million bp) and low scaffold contiguity around these genes for *D. persimilis* and *D. hydei*.

### ChIP-seq, RNA-seq and ATAC-seq analyses

Raw data of genome binding/occupancy (ChIP-seq), transcriptome (RNA-seq) and nucleosome (ATAC-seq) profiling were obtained from GEO database and used in the analyses. They include: GSE59965 - contains data for *D. virilis* including RNA-seq, ChIP-seq of H3K9me3 and RNA polymerase II performed using commercially available anti-H3K9me3 (ab8898, Abcam) and anti-RNA Pol II (ab5408, Abcam) antibodies; GSE35648 - contains data for both *D. melanogaster* and *D. virilis* including ChIP of BEAF-32 performed using antibodies generated against amino acids 1-83 of the major highly conserved isoform BEAF-32B in *D. melanogaster* (*90*); GSE43829 - contains RNA-seq as well as ChIP-seq of H3K9me3 and RNA polymerase II for *D. melanogaster* performed using aforementioned commercially available antibodies ab8898 and ab5408; GSE56347 - includes ChIP-seq of HP1a for *D. melanogaster* performed with polyclonal anti-HP1 (PRB-291C, Covance innovative); GSE102439 - includes ATAC-seq data for *D. melanogaster* and *D. virilis*; finally, GSE85404, GSE33052, GSE70632 and GSE76997 - contains ChIP-seq data of dCTCF, Su(Hw), GAF, Pita, ZIPIC and Zw5 for *D. melanogaster*.

For analysis of sequence data, we used genome sequence and annotations released in FlyBase, *D. melanogaster* r.6.19 and *D. virilis* r1.06. Prior to mapping, all libraries were subjected to adapter clipping, filtering by length (>20 nt) and quality (80% of nt must have at least 20 Phred quality) using TrimGalore (https://github.com/FelixKrueger/TrimGalore). Then, sequences were aligned to corresponding genomes using Bowtie (91) with the following settings: “-v 0 --best --strata”, retaining only aligned reads with zero mismatches. Output sequence alignment map (SAM) files were converted to binary (BAM) format using SAMtools (92). BAM files were converted to wiggle (WIG) format using bamToWig script (https://github.com/craiglowe/bamToWig). Aligned reads in wig format were visualized using the Integrative Genome Viewer (IGV) (93).

Peak calling was performed using MACS software (94) with the recommended parameters for narrow (PolII, BEAF-32, dCTCF, GAF, Su(HW), Pita, ZIPIC, Zw5) and broad peak calling (H3K9me3, HP1a). Enrichment analysis was performed using pipelines implemented in deepTools package (95) with the parameters including ignoring of duplicates.

For ATAC-seq analysis, reads that mapped on mitochondrial genomes were discarded, and peak calling was performed using Genrich (https://github.com/jsh58/Genrich) with the following settings: “-j -y -r -d 50”, including removal of PCR duplicates.

### Promoter analysis

Because of insertion of DAIBAM MITE at the distance of 92 bp upstream from TSS of *Myb* in *D. virilis*, the promoter regions of *Myb* in studied species were shortened to 100 bp. After sequence extraction, promoter regions of *Myb* and *Ranbp16* were searched for common motifs using MEME (96) and identification of matches to known transcription factors was performed by Tomtom (97) using OnTheFly (39) and REDfly v5.6 (40, 41) databases implemented in MEME Suite 5.0.5 (38).

### Sequence evolution and testing for selection

Analysis of nucleotide substitutions per site was conducted in MEGA X (98) using the Tamura-Nei model (99). Rate variation among sites was modeled with a gamma distribution (shape parameter = 1). All positions containing gaps and missing data were eliminated (complete deletion option).

Ratio of nonsynonymous and synonymous substitutions (dN/dS) was estimated using PAL2NAL software (100) by converting multiple sequence alignment of proteins and the corresponding nucleotide sequences into a codon alignment, and the calculation of synonymous (dS) and non-synonymous (dN) substitution rates using codeml program implemented in PAML package (101).

### Cytology and DNA in situ hybridization

*D. virilis* larvae were grown at 18^0^C on standard medium supplemented with live yeast solution for 2 days before dissection. Salivary glands from 3^rd^ instar larvae were dissected in 45% acetic acid and squashed. DNA probes corresponding to *D. virilis Myb* (Dvir\GJ18431; FlyBase ID: FBgn0205590) were prepared by PCR using gene-specific primers (Forward_ GCAAGTGCGAGCACTGAAAA; Reverse_TGCATACTGAGGTGTGCCAG). Then, DNA probe was biotinylated by nick translation using Biotin-14-dATP (Thermo Fisher Scientific, USA) as described in (102). Localization of the probe was made using the cytological map of *D. virilis* chromosomes (103). Images were obtained by binocular microscope Nikon Alphaphot-2 YS2 (Japan).

### RNA isolation, RT-PCR and 5’-RACE analysis

Total RNA from 3^rd^ instar larvae, adult females and gonads was isolated using Extract RNA reagent (Evrogen, Russia). Synthesis of the first strand of cDNA from total RNA and subsequent amplification of regions of interest were performed using MINT cDNA kit (Evrogen, Russia) following manufacturer’s instructions. For specific rapid amplification of cDNA 5’-end (5’-RACE) analysis, we applied two outward primers (primer1 5’-AGTAGTTGTGCGTAGCTGGA-3’; primer2 5’-GCTGCTTGCACAATGTTTCTA-3’) corresponding to the annotated 5’-fragment of *D. virilis Ranbp16* gene (Dvir\GJ18467; FlyBase ID: FBgn0205626). PCR reaction was conducted using Encyclo DNA polymerase (Evrogen, Russia). The resulting PCR fragments were cloned into pAL2-T vector (Evrogen, Russia) and sequenced using plasmid-specific primers. In all RT-PCR experiments, probes containing all components but lacking reverse transcriptase were used as negative controls. The obtained sequence of 5’UTR of *D. virilis Ranbp16* gene was deposited in GenBank under the number MN481598.

## ACCESSION NUMBERS

The sequence of 5’UTR of *D. virilis Ranbp16* gene was deposited in GenBank under the number MN481598.

## Financial disclosure

This work was supported by the Russian Foundation for Basic Research (grant number 19-04-00337). The funders had no role in study design, data collection and analysis, decision to publish, or preparation of the manuscript.

## Acknowledgements

We are grateful to Dr. Anton Golovnin from the Institute of gene biology RAS for ideas and critical comments on the manuscript. We thank Alexei M. Kulikov from the Koltzov Institute of developmental biology RAS for helpful advices and technical assistance. The bioinformatics was performed using the computational facilities of Engelhardt Institute of Molecular Biology RAS Genome center (http://www.eimb.ru/rus/ckp/ccu_genome_c.php).

## Supporting Information Legends

**Supplementary figure S1.** The content of repetitive DNA in the genomic region of *D. virilis* comprising *Myb* and *Ranbp16* genes. Classes and subclasses of transposable elements are indicated in different colors. Divergence of repeats is shown in the percentage of element coverage of the entire length of the consensus sequence where 100% should be considered as entire coverage of repeat consensus sequence on genome region.

**Supplementary figure S2.** The estimated evolutionary divergence (A and B) and the ratio of non-synonymous and synonymous substitutions (C and D) across Diptera species. A and B) The number of base substitutions per site (Distance) for *Myb* and *Ranbp16* genes, respectively. Standard error estimate(s) are shown above the diagonal (Standard error). Analyses were conducted using the Tamura-Nei model. All positions containing gaps were discarded. C and D) Results of dN/dS test for *Myb* and *Ranbp16* genes, respectively. Estimation was performed in relation to *D. melanogaster* sequence.

**Supplementary figure S3.** Multiple protein alignment of Myb type HTH DNA-binding domain of Myb (A) and Importin-β domain of Ranbp16 (B) across selected taxa.

**Supplementary figure S4.** The intergenic region between *Ranbp16* gene and the adjacent *Stim* gene of *D. melanogaster* exported from UCSC genome browser. CAGE-seq data indicate the beginning of 5’UTR (5’untranslated region) on each DNA strand. Top upper track shows the promoter region coordinates fetched from Eukaryotic Promoter Database (EPD) (https://epd.epfl.ch//index.php). Sequence conservation is shown for 27 insects presented in the UCSC database.

**Supplementary figure S5.** The enrichment profile of insulator proteins in the heterochromatin of *D. melanogaster*. A) The enrichment profile of insulators BEAF-32, dCTCF, GAF, Pita, ZIPIC, Su(Hw), Zw5 and heterochromatic marks H3K9me3 and HP1a in the pericentric region of chromosome 2L of *D. melanogaster*. The euchromatin-heterochromatin border is depicted by dashed line according to the mapping profile of H3K9me3 and HP1a. Mapped ChIP-seq reads without normalization (mapped reads) and calculated areas of enrichment relative to the input data with the *p*- and *q*-values < 0.05 (peaks) are shown for each factor. RNA-seq reads were normalized to the sequence depth (RPM, reads per million). B) The enrichment profile of insulator proteins and H3K9me3 upstream, within and downstream of heterochromatic genes located in pericentric region of chromosome 2L of *D. melanogaster*.

**Supplementary table S1.** List of the *Myb* and *Ranbp16* orthologs and their coordinates in the genome.

**Supplementary table S2.** List of the studied heterochromatic genes of *D. melanogaster* and their orthologs in *D. virilis* with coordinates in the genome.

